# The metabolic profile of lag and exponential phases of *Saccharomyces cerevisiae* growth changes continuously

**DOI:** 10.1101/063909

**Authors:** Rafael N. Bento, Miguel A. Aradhya, Valdir A. R. Semedo, Carlos E. S. Bernardes, Manuel E. M. Piedade, Fernando Antunes

**Affiliations:** Centro de Química e Bioquímica e Departamento de Qumica e Bioquímica, Faculdade de Ciências, Universidade de Lisboa, 1749-016 Lisboa, Portugal.

**Keywords:** flow-microcalorimetry, exponential phase, lag phase, metabolic profile, oxidative respiration

## Abstract

Cellular growth is usually separated in well-defined phases. For microorganism like *Saccharomyces cerevisiae,* two phases usually defined are (1) a lag phase, in which no growth is observed and cells adapt to a new environment, followed by (2) an exponential phase, in which rapid proliferation occurs. Here we investigate whether these well-defined phases are uniform. By using flow-microcalorimetry, we found that the metabolic profile of the culture is continuously changing, both in the lag and exponential phases of growth. Along the lag phase there is a continuous increase in the energy that is dissipated irreversibly as heat, while in the exponential phase the opposite occurs. We also confirm recent observations that the oxidative component of metabolism decreases along the exponential phase. Interestingly, nutrient limitation further decreases the amount of energy that is dissipated irreversibly. Altogether, this points to a picture in which cells respond rapidly to minute environmental changes by adjusting their metabolic profile.

## 1 INTRODUCTION

Understanding how cells use nutrients to promote cellular growth and maintenance is a fundamental biological problem that has been under intense investigation. Anabolic reactions needed for biosynthetic processes require an input of free energy in the form of ATP that is harnessed from nutrients in metabolic pathways such as glycolysis and mitochondrial respiration. Mitochondrial respiration generates 36 molecules of ATP per molecule of glucose while glycolysis generates two. Thus, it is puzzling that a high rate of glycolysis in the presence of oxygen (aerobic glycolysis), in which glucose is only partially oxidized, for example to lactate or ethanol, rather than being oxidized to CO_2_ in respiration, is observed in many situations. Known as the Warburg effect, following the observation of aerobic glycolysis by Otto Warburg in cancer cells near a century ago, this phenomenon is also observed in non-cancer fast-proliferating cells such as lymphocytes [1] [2] during development [3], stem cell proliferation [4], lung fibrosis [5], in many unicellular organisms exposed to a sugar-rich environment, such as *Saccharomyces cerevisiae* (Sc). Recently, it was found that the exponential growth phase of Sc, in which there is a rapid cell division, is not an uniform phase, but instead, shows a highly dynamic metabolism being more oxidative in the early exponential phase and more fermentative at the end of the exponential phase [6]. This was surprising because the time for cell division is constant along the exponential phase, and previous observations indicated that metabolic fluxes during the exponential phase are in a near steady-state. The evidence suggesting the dynamic nature of the exponential phase was very robust resulting from a flux analysis of CO_2_ production, O_2_ consumption and ATP generation, and included also an evaluation of stress resistance and a genomic and proteomic profiling along the exponential phase 6. In addition, previous observations that substrate consumption and product formation per unit of biomass formed varies during different time periods within the exponential phase also revealed a metabolic variation along the exponential phase [7].

In this work, we analyzed the issue of metabolic uniformity over the exponential phase by following Sc growth with calorimetry, which is the gold standard to measure metabolic rates [7]. Being a non-invasive, probe-free and extremely sensitive technique, calorimetry measures metabolic activity in real time as the heat released by organisms or cells [8][9]. This heat represents the free energy dissipated irreversibly during metabolism, providing an all-inclusive measure of metabolic activity. We built up on previous studies on yeast microcalorimetry [7][10][11], and by applying state of the art electronics for data acquisition and analysis, highly accurate reproductive data were obtained with our flow microcalorimetry set-up. As opposed to previous studies in which prototrophic strains were used [7] [6], we analyze an auxotrophic strain (BY4741) that is commonly used in laboratory studies [12]. Main observation was the non-uniform and highly-dynamic nature of the exponential growth phase, with the dissipated heat together with the rate of respiration decreasing along the exponential phase. The maximal dissipated heat occurs at the transition between the lag to the exponential phase. In conclusion, we confirm and extend previous observations that metabolic changes occurring during the exponential phase favor aerobic glycolysis over mitochondrial respiration.

## 2 Materials and Methods

### 2.1 Yeast strain, media, and growth conditions

*Materials*— This work was performed with the *Saccharomyces cerevisiae* (Sc) haploid strain BY4741 (wild-type, genotype MATa his3Δ1 leu2Δ0 met15Δ0 ura3Δ0), obtained from EUROSCARF (Frankfurt, Germany). Yeast extract, bactopeptone, yeast nitrogen base and agar were from Becton, Dickinson and Company (Le Pont de Claix, France). Glucose was obtained from Merck, (Darmstadt, Germany). The amino acids and nitrogen bases were from Sigma Chemical Company (St. Louis, MO, USA).

*Media and growth conditions*—Growth and all incubations were made in either Yeast Peptone Dextrose (YPD) containing 1% (w/v) yeast extract, 2% (w/v) peptone and 2% (w/v) D-glucose, or in Synthetic Complete (SC) medium containing 0.685% (w/v) yeast nitrogen base, 2% (w/v) D-glucose and the amino acids and nitrogen bases as indicated in [13] at 30 °C, under aerobic and batch conditions and shaking at 160 rpm in an Infors AG CH-4103 Bottmingen incubator or under vigorous shaking in an incubator next to the calorimeter. Medium was always prepared one day before each experiment because this procedure led to a higher reproducibility in comparison with older media.

### 2.2 Statistical analysis

Results are presented as the means and the assigned uncertainties correspond to standard deviations of the number *n* of independent determination given in parenthesis. Data statistical analysis was undertaken using a two-tailed Student *t* test; differences were significant when the p-value was lower than 0.05 (* - p < 0.05; ** - p < 0.01).

### 2.3 Calorimetry

Calorimetric experiments were carried out with a previously described [14] LKB 10700-1 flow calorimeter operated in the flow-through mode. The original apparatus was, however, expanded with ancillary equipment for cell growth. The cell cultures were kept in a jacketed glass vessel that was placed inside an incubator adjacent to the calorimeter. Magnetic stirring ensured an appropriate oxygenation of the culture media and constant humidity was achieved by placing an open vessel filled with distilled water inside the incubator. The cell cultures were circulated between the glass vessel and the calorimetric cell inside Teflon tubes. Pumping was by means of an Ismatec MS-4/12 multi-channel peristaltic pump located inside the incubator. The whole system was thermostated at 30°C, including the Teflon tubes used to transport the cell culture between the glass vessel and the calorimetric cell. This last feature was achieved by keeping the Teflon lines exiting the incubator and returning from the calorimeter inside flexible PVC pipe bellows, where thermostated air from the incubator was circulated by a system of fans. Different thermostatic units ensured a temperature stability of ±0.01 K in the glass vessel (HAAKE DC 5 unit), ±0.03 K in the incubator (Julabo LC6 controller), and ±0.008 K in the calorimeter (LKB 10700-1 air thermostat jacket). The thermostatization of the glass vessel inside an already temperature controlled environment allowed a better stabilization of the cell culture temperature, which became immune to the small variations associated with the opening/closing of the incubator during the experiments. The all apparatus was kept in an air-conditioned room whose temperature was regulated to 22±1ºC.

The culture volume in the reaction vessel was 20 mL. The nominal volume flow rate in the peristaltic pump was adjusted so that an actual flow rate *q_V_* = 1.0 mL.min^-1^ was achieved in all experiments. This adjustment was regularly checked as flow rate variations affect various calorimetric parameters, such as the calibration constant [14], the effective or thermal volume of the calorimetric vessel [14] or the signal sensitivity [15].

The flow calorimetric line and the calorimetric vessel were sterilized after each experiment by circulating: (i) an aqueous solution of bleach 20 % (v/v) for, at least, 4-5 h; (ii) deionized water from a millipore system for 10-20 min; (iii) aqueous ethanol 70 % (v/v) for 20 min; and (iv) millipore H_2_O for 10-20 min. The reaction vessel and other glassware were sterilized at 120°C during 2.5 h. Alternatively the system was also sterilized by circulating ethanol 70 % (v/v) for at least 3 h and then circulating H_2_O Millipore for 1 h to wash cellular debris.

Electrical calibrations of the calorimetric system under the same conditions used in the main experiments (e.g. the pump flow rate, temperature) were performed as previously described [14]. No significant differences in the calibration constant, *ɛ*, were observed when millipore water, SC media or a culture of Sc in stationary growth phase were flown through the calorimetric cell. The use of cells in stationary phase allowed the recording of a stable baseline during an extended period of time. The value *ɛ* = 15.5±0.2 μ.WμV^-1^ (*n* = 41) was used throughout this work.

In order to assign the power dissipated by a cell culture to a specific cell number or biomass it is necessary to know the thermal or effective volume of the calorimetric cell [16], which differs from its physical volume since the flow of cell culture results in transport of heat away from the detection area. The effective volume of the LKB 10700-1 flow-through cell, *V*_eff_ = 0.547±0.012 mL (n=4) under the temperature and volume flow rate conditions of the present experiments was determined as recommended in the literature [16], using the base catalyzed hydrolysis of methyl paraben. The calculation of *V* _eff_ relied, however, on the standard molar enthalpy of that reaction at 30°C, Δ_r_ 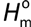 = - 50.60±0.82 kJ-mol^-1^ (unpublished results) recently obtained in our laboratory, which is more precise than the previously reported benchmark [16].

Experiment and calibration programing, and data acquisition, were controlled with the CBCAL 1.0 software [17].

### 2.4 Experimental procedure

A pre-culture of Sc cells at stationary phase was grown overnight and then inoculated at an initial OD_600_ ~0.035, where OD_600_ represents the optical density at 600 nm. The cell culture was divided in two and the resulting preparations were followed in parallel: one (A) calorimetrically and the other (B) through OD_600_ measurements every hour. Culture A was transferred to the calorimetric apparatus immediately after inoculation. Culture B was grown in an Erlenmeyer flask under magnetic stirring. The flask was either placed inside the incubator adjacent to the calorimeter, that also contained culture A, or in another incubator (AG CH-4103 Bottmingen) that was set to 30°C. To ensure that parallel growth is indeed observed for cultures followed through OD_600_ measurements and calorimetry, the OD_600_ of culture A was measured at the end of every experiment and compared with that of culture B.

### 2.5 Biomass and cell number

Biomass was determined using a calibration curve plotting OD_600_ to dry weight biomass. For dry weight determinations, cells were grown in SC media as described above and 50 mL samples were taken every hour, centrifuged at 5000 g for 5 min at 4 °C in a Eppendorf centrifuge 5804 R, washed once with distilled water, centrifuged again and stored at −20 °C until lyophilisation, carried out for at least 24h. Samples were then weighted with a precision of 0.01 mg using a Mettler Toledo XS205 balance.

The unit carbon formula weight considered was 24.4 g_biomass_ C-mol^-1^ 7. Although this value is not specific for the BY4741 strain is a close approximation since the elemental composition between different strains of yeast is fairly constant [18].

Conversion of OD_600_ to cell number was calculated from plots of OD_600_ versus cells/mL plots. A cell culture growth was accompanied by OD_600_ and by counting cells in a hemocytometer. A value of 1 OD_600_ = (2.94±0.12)x10^7^ cells.mL^-1^ was obtained as the mean of four slopes (*n* = 4).

### 2.5 Biochemical determinations

The oxygen consumption rate was measured by transferring 0.8 mL of a growing culture of Sc into a *Hansatech Oxygraph plus* (Hansatech Instruments Ltd., Norfolk, UK) thermostatically controlled at 30°C. Calibration was made using a N_2_ flush to the 0% O_2_ signal and a compressed air flush until the signal saturated at 21 % O_2_.

Glucose concentration was determined by transferring 0.8 mL of medium (at the required dilution, if necessary) into a *Hansatech Oxygraph plus* at 30°C and then, when baseline was stable, 20μL of glucose oxidase (14.5 U) (Sigma Chemical Company, St. Louis, MO, USA) were added with a Hamilton syringe. The glucose oxidase will specifically oxidize the D-glucose in the medium, consuming oxygen in the process, which is quantified by the electrode. A calibration curve was used to convert the output signal into glucose concentration.

## 3 Results and Discussion

### 3.1 Growth in synthetic complete medium

To test whether the exponential phase of growth is uniform, we started by monitoring growth of Sc cells in SC medium. Figure 1A shows the results of two parallel experiments where, cell growth was followed by optical density and calorimetric measurements. The power (*P)* derived from the calorimetric signal is a measure of the heat produced by the overall cell population per unit time. The observed *P-t* profile is similar to that observed by others [11][10] and starts with the lag phase, in which microorganisms adapt to a new rich medium and divide very slowly. A low calorimetric signal is detected during this phase. After the adaptation period, the power rapidly increases, corresponding to the exponential growth phase, in which a rapid division of cells is driven by a respiro-fermentative metabolism, with the number of cells showing a generation time of 0.44±0.03 h^-1^ (*n* = 26).

**Figure 1.**
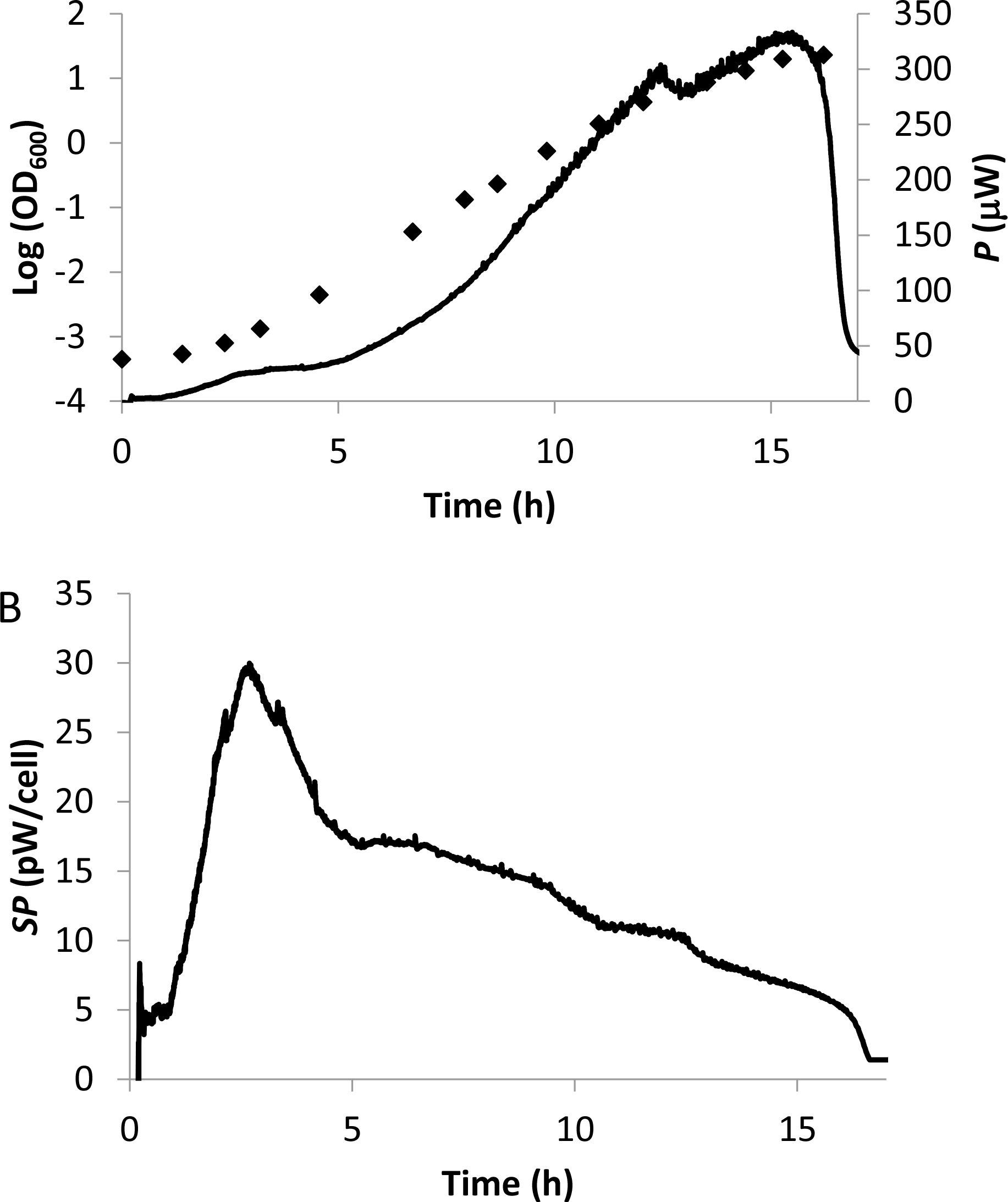
Growth curve of Sc in SC medium. In (A) logarithm of optical density at 600 nm (♦) and dissipated power (*P*) (―――――――) are plotted. In (B) the specific power (*SP*) (―――――――) refers to the ratio of *P* (shown in panel 1A) over the number of cells.

Additional information is obtained from the specific power, *SP,* which is calculated by dividing *P* by the number of cells. The *SP-t* plot (Figure 1B) considerably differs from the *P-t* plot and gives important insights on the metabolic dynamics along the Sc growth cycle. One important aspect of this curve is that the maximum value of *SP,* 26±3 pW cell^-1^ (*n* = 6), is observed at the end of the lag phase. In other words, as suggested before [19], the lag phase, where cell division is slow, is characterized by the most intense dissipative metabolism within the growth cycle. After the lag phase, there is an abrupt decrease in *SP,* and then *SP* values continuously drop along exponential growth, never stabilizing. There is a large variation, from *SP* = 20±2 pW cell^-1^ (*n* = 5) at the beginning to *SP* = 5±1 pW cell^-1^ (*n* = 5) at the end of the exponential phase. This indicates that exponential growth does not correspond to a steady uniform phase, but instead is highly dynamic and includes continuous metabolic changes.

Because the enthalpy associated with full oxidation of glucose to CO_2_ is higher than that associated to oxidation of glucose to ethanol, i.e. fermentative metabolism is less dissipative than respiration [18][20], a possible explanation for the observed decrease of *SP* along the exponential phase is a decrease of the respiratory metabolism, as observed in [6]. Figure 2 shows that a decrease of the rate of consumption of O_2_ along the exponential phase is indeed observed, i.e., oxidative metabolism decreases resulting in a decrease of the glucose fraction that is fully oxidized to CO_2_ along the exponential phase. This further supports the conclusion that respiration contribution to metabolism decreases along the exponential phase.

**Figure 2.**
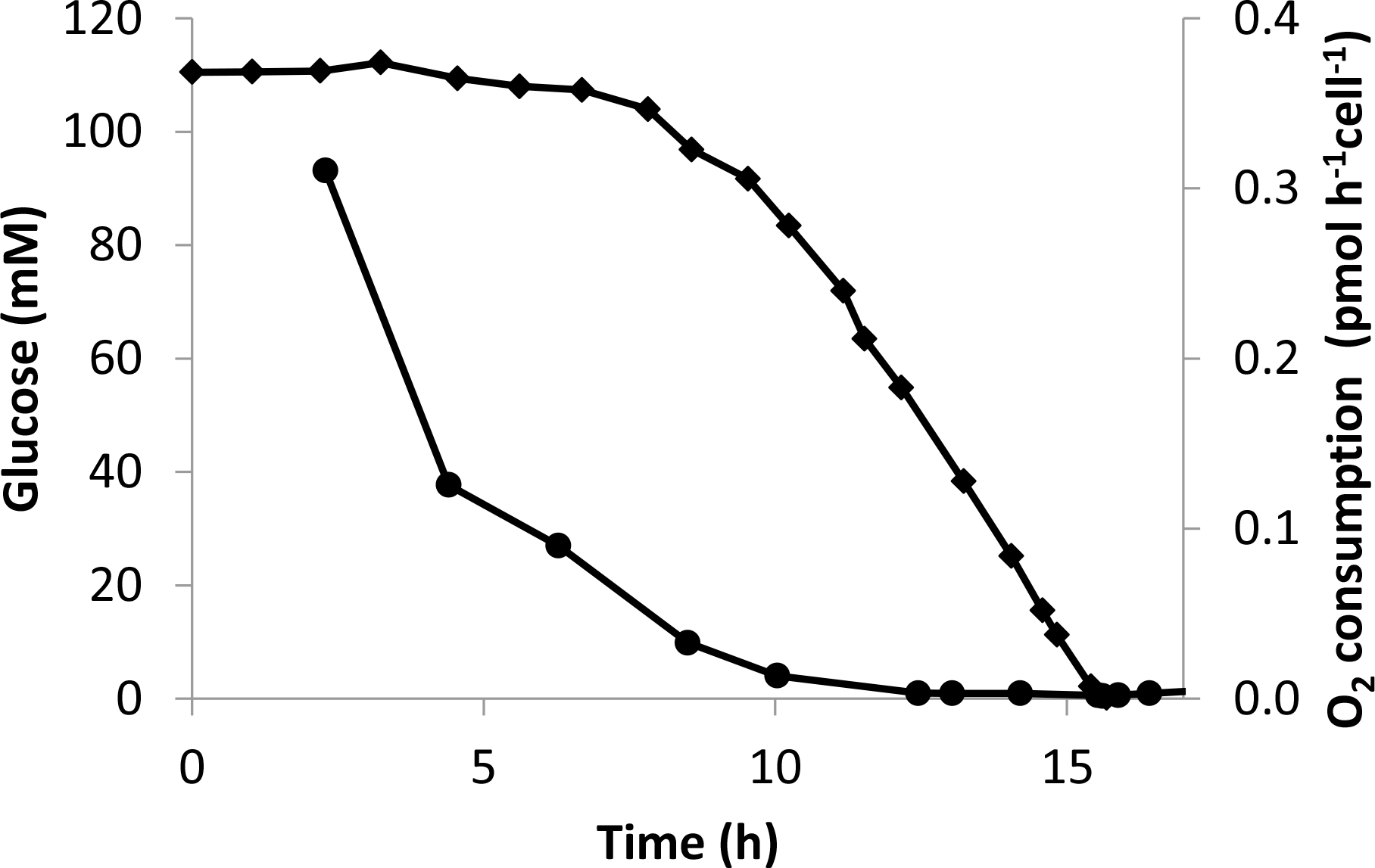
Glucose culture levels (♦) and rate of O2 consumption (•) during the growth of Sc cells in SC medium.

The reasons for the dynamic character of the exponential phase are not clear. One possible explanation is that by switching to a more fermentative metabolism Sc cells have the ability to maintain a constant cell division rate. Nevertheless, the period in which a constant cell division rate is observed cannot be extended beyond 10.5-11 h, when a drop in the duplication rate to 0.17±0.02 h^-1^ (*n* = 5) is observed (Figure 1A). It should be mentioned that during this slower growth phase, there is still plenty of glucose available (approximately 60 mM) and the metabolism continues to be mostly fermentative with little O_2_ consumption (Figure 2) dissipating even less energy per cell than in the first 10 h of growth (Figure 1B). This behavior reveals that SC medium is not able to support an exponential fast-dividing phase while glucose is still present, an observation that may result from the auxotrophy of this strain [12] or limiting inositol levels in SC medium [21]. In fact, the biomass yield per C-mol of glucose at the beginning of the growth cycle is 0.264±0.023 C-mol_biomass_/C-mol_g_|_ucose_ (*n* = 5) while at the end it decreases by near 50 % to 0.125±0.008 C-mol_biomass_/C-mol_glucose_ (*n* = 5). This decrease in the biomass yield per C-mol of glucose consumed is consistent with the glucose wasting observed when auxotrophic nutrients are limiting the growth [22] [23].

### 3.2 Growth in a rich medium

Due to the sensitivity of BY4741 to the media used [12] [24], we evaluated if the type of medium could affect the dynamics of metabolism change during the exponential phase by growing Sc cells in YPD medium, which is richer than SC, thus supporting higher proliferation rates. Optical density and *P-t* measurements (Figure 3A) give experimental profiles similar to those observed in Figure 1A. As expected the generation time 0.52±0.02 h^-1^ *(*n** = 3) was higher than that obtained with the poorer SC medium. Concerning the *SP* plot (Figure 3B), the highest *SP* value, approximately 34±2 pW cell^-1^ (*n* = 3), was observed at the end of the lag phase, similar to what was noted for cells growing in SC medium. Remarkably, during the exponential phase the results show a similar behavior to that observed for the growth curve obtained in SC medium. During the exponential phase, after the initial abrupt decrease following the lag phase, *SP* values also drop continuously never stabilizing. There is a large variation, from 26.2±2.2 pW cell^-1^ (*n*=2) at the beginning to 6.4±0.6 pW cell^-1^ (*n* = 3) at the end of the exponential phase. This indicates that the non-uniformity of this period of growth is not dependent on the type of growth medium used.

**Figure 3.**
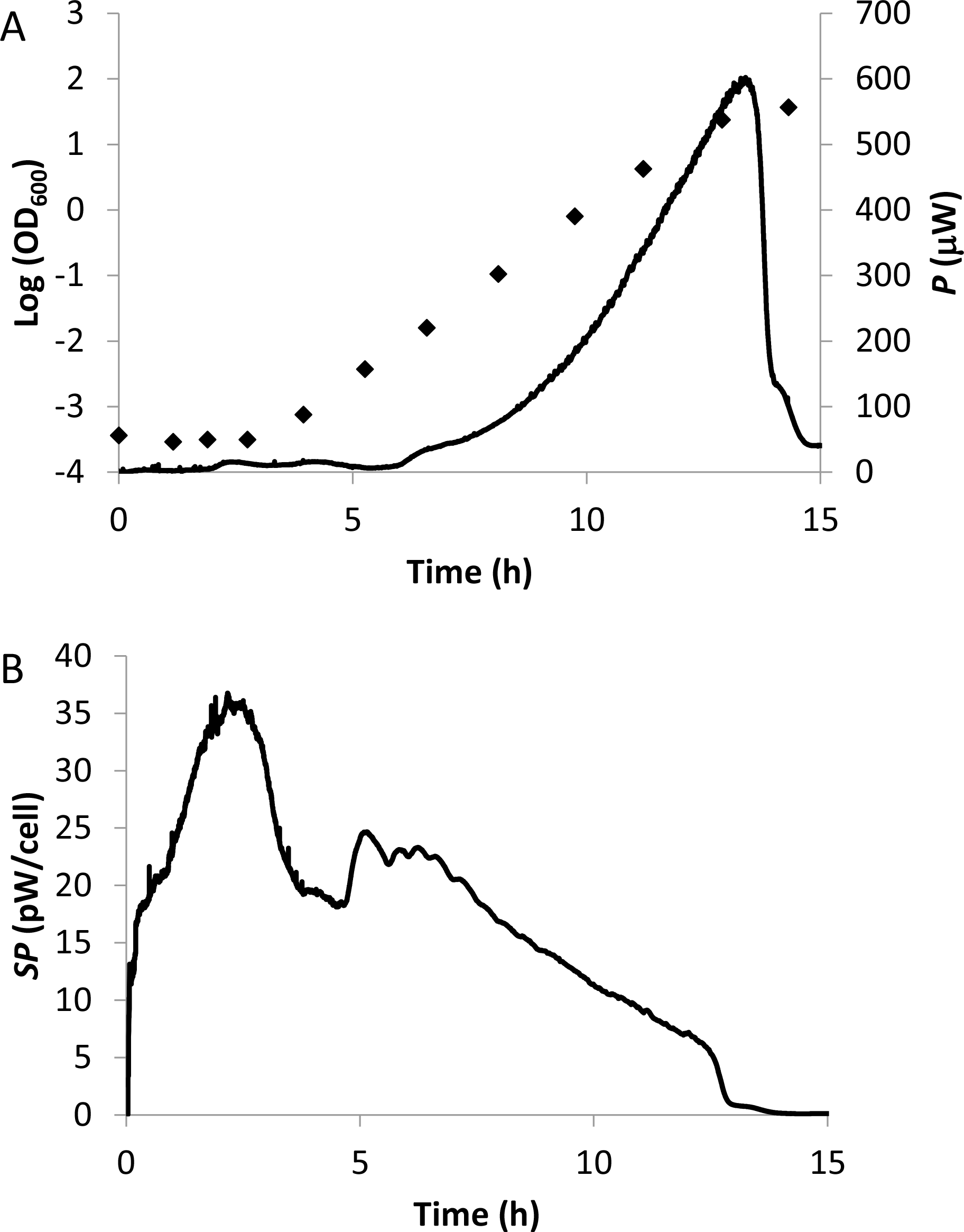
Growth curve of Sc in YPD medium. In (A) logarithm of optical density at 600 nm (♦) and dissipated power (*P*) (―――――――) are plotted. In (B) the specific power (*SP*) (―――――――) refers to the ratio of *P* (shown in panel 1A) over the number of cells.

### 3.3 Final remarks

Several mechanisms contribute to the occurrence of aerobic glycolysis. The decrease in the power dissipated per cell observed during the exponential phase is an indication that aerobic glycolysis may spare nutrients to allow a fast proliferation rate. This is further supported by the observation that in the poorer SC medium, when nutrients are not able to support a full proliferation rate but glucose is still present at high levels, the dissipated heat per cell further decreases, indicating an even lower contribution of respiration to cell metabolism than in the exponential phase. Aerobic glycolysis is not associated only to fast cell proliferation. In fact, aerobic glycolysis has also been observed in situations where growth is limited by the lack of nutrients, namely auxotrophic nutrients [22][23]. The dynamic nature of the exponential phase is probably not a universal phenomenon, as many studies show that this phase can be considered a true uniform period of growth with metabolism near a steady-state. Strain difference, among other experimental conditions, may account for these discrepancies, and it will be desirable if a common strain is used as a benchmark to facilitate comparison of data between studies allowing a faster development of yeast research, as proposed by van Dijken *et al.* [25]. Altogether, the high sensitivity of calorimetry measurements can give important insights on the metabolic profile of cells, contributing to a better understanding of how cells use nutrients to promote cellular growth and maintenance, and respond to excess or limitation of some nutrients, issues that are at the center of many metabolic diseases.

## Acknowledgments

This work was supported by FCT, Portugal, through Project PEst-OE/QUI/UI0612/2013 and a postdoctoral grant awarded to C. E. S. Bernardes (SFRH/BPD/101505/2014). We acknowledge Hermínio P. Diogo for its contribution to the design of the reaction vessels.

